# Cinnamon bark extract suppresses metastatic dissemination of cancer cells through inhibition of glycolytic metabolism

**DOI:** 10.1101/2021.03.25.437098

**Authors:** Yuki Konno, Ami Maruyama, Masaru Tomita, Hideki Makinoshima, Joji Nakayama

## Abstract

Metastasis, a leading contributor to the morbidity of cancer patients, occurs through multiple steps. As each of these steps is promoted by different molecular mechanisms, blocking metastasis needs to target each of these steps. Here we report that cinnamon bark extract (CBE) has a suppressor effect on metastatic dissemination of cancer cells. Though a zebrafish embryo screen which utilizes conserved mechanisms between metastasis and zebrafish gastrulation for identifying anti-metastasis drugs, CBE was identified to interfere with gastrulation progression of zebrafish. A zebrafish xenotransplantation model of metastasis validated that CBE suppressed metastatic dissemination of human cancer cells (MDA-MB-231). Interestingly, quantitative metabolome analyses revealed that CBE-treated MDA-MB-231 cells disrupted the production of glucose 6-phosphate (G6P) and fructose 6-phosphate (F6P), which are intermediate metabolites of glycolytic metabolism. CBE decreased the expression of hexokinase 2 (HK2), which catalyzes G6P production, and pharmacological inhibition of HK2 suppressed cell invasion and migration of MDA-MB-231 cells. Taken together, CBE suppressed metastatic dissemination of human cancer cells by inhibiting glycolytic metabolism.

## Introduction

Metastasis substantially contributes to morbidity and mortality in cancer patients. Its progression involves multiple steps: invasion, intravasation, survival in the circulatory system, extravasation, colonization, and secondary tumor formation in new organs accompanied by angiogenesis [1]. The physical translocation of cancer cells is an initial step of metastasis and molecular mechanisms of it involve cell motility, breakdown of the local basement membrane, loss of cell polarity, acquisition of stem cell-like properties, and epithelial-to-mesenchymal transition. These cellular phenomena resemble the evolutionarily conserved morphogenetic movements observed during vertebrate gastrulation, including epiboly, internalization, convergence, and extension [2].

The zebrafish system has been increasingly recognized as a platform for chemical screening because it provides the advantage of high-throughput screening in an *in vivo* vertebrate setting with physiological relevance to humans [3, 4]. We previously developed a screening concept utilizing conserved mechanisms between metastasis and zebrafish gastrulation for identifying anti-metastasis drugs. The screen used epiboly, the first morphogenetic movement in gastrulation, as a marker and enabled 100 chemicals to be tested in five hours [5].

Cinnamon bark (CB) is one of the most popular spices obtained from the inner bark of several tree species from the genus *Cinnamomum*. CB extract (CBE) contains various bioactive compounds: cinnamic aldehyde, cinnamic acid, carbohydrates, tannins, and mucus. Previous studies found that these compounds have antidiabetic, antioxidative, anti-inflammatory, and anticancer effects [6–8].

Here we show that CBE suppresses metastatic dissemination of human cancer cells through inhibition of glycolytic metabolism.

## Materials and Methods

### Zebrafish embryo screening

Each herbal medicine was added at a final concentration of 100 μg/ml to a well of a 24-well plate containing approximately 20 zebrafish embryos at the sphere stage per well. The screening assay was performed as described previously [5]. All experimental zebrafish were handled in accordance with institutional guidelines established by the Animal Care Committee of the National Cancer Center.

### Reagents

The herbal medicine library and CBE were provided by Prof. Hayakawa, Institute of Natural Medicine, University of Toyama, Toyama, Japan.

### Cell culture and cell viability assay

MDA-MB-231 cells were obtained from the American Type Culture Collection (ATCC, Manassas, VA).

### Boyden chamber cell motility and invasion assay

The cell motility and invasion assays were performed as previously described [9]. In the Boyden chamber assay, aliquots containing 3 × 10^5^ MDA-MB-231 cells were dispensed to each well in the upper chamber.

### Immunoblotting

Western blotting was performed as described previously [10]. Anti-HK2 and GAPDH antibodies are purchased from Cell Signaling Technology.

### Zebrafish xenotransplantation model

*Tg (kdrl: eGFP)* zebrafish was provided by Dr. Stainier (Max Planck Institute for Heart and Lung Research). Embryos that were derived from the line were maintained in E3 medium containing 200 μM 1-phenyl-2-thiourea (PTU). Approximately 100–400 red fluorescence protein (RFP)-labeled MBA-MB-231 cells were injected into the duct of Cuvier of the zebrafish at 2 days post-fertilization (dpf).

### Measurement of extracellular acidification rate (ECAR)

ECAR was measured using the XF glycolysis stress test kit according to the manufacturer’s instructions (Seahorse Bioscience, Agilent).

### Metabolite measurements

Metabolic extracts were prepared from 1–5 × 10^6^ MDA-MB-231 cells in methanol containing an internal standard solution and subjected to metabolite analyses using a gas chromatography-mass spectrometry (GC-MS) system.

### Statistics

Data were analyzed using Student’s t test, and p < 0.05 was considered statistically significant.

## Results

### Zebrafish embryo screen identified CBE as an epiboly-interrupting herbal medicine

A previous study demonstrated that chemicals that interfere with the epiboly progression in zebrafish embryos could suppress metastasis [5]. To identify the herbal medicine that has suppressor effects on metastasis, we subjected 162 Herbal medicines into the zebrafish embryo screen. We found that 14.8% (24/162) of the herbal medicines either delayed or interrupted epiboly of the embryos. Among these 24 medicines, CBE severely delayed embryonic cell movement. No effect on the epiboly progression of the embryos was detected for 70% (114/162) of the herbal medicines, whereas 12% (20/162) of the medicines induced toxic lethality (Figure 1A and B).

**Figure 1.**
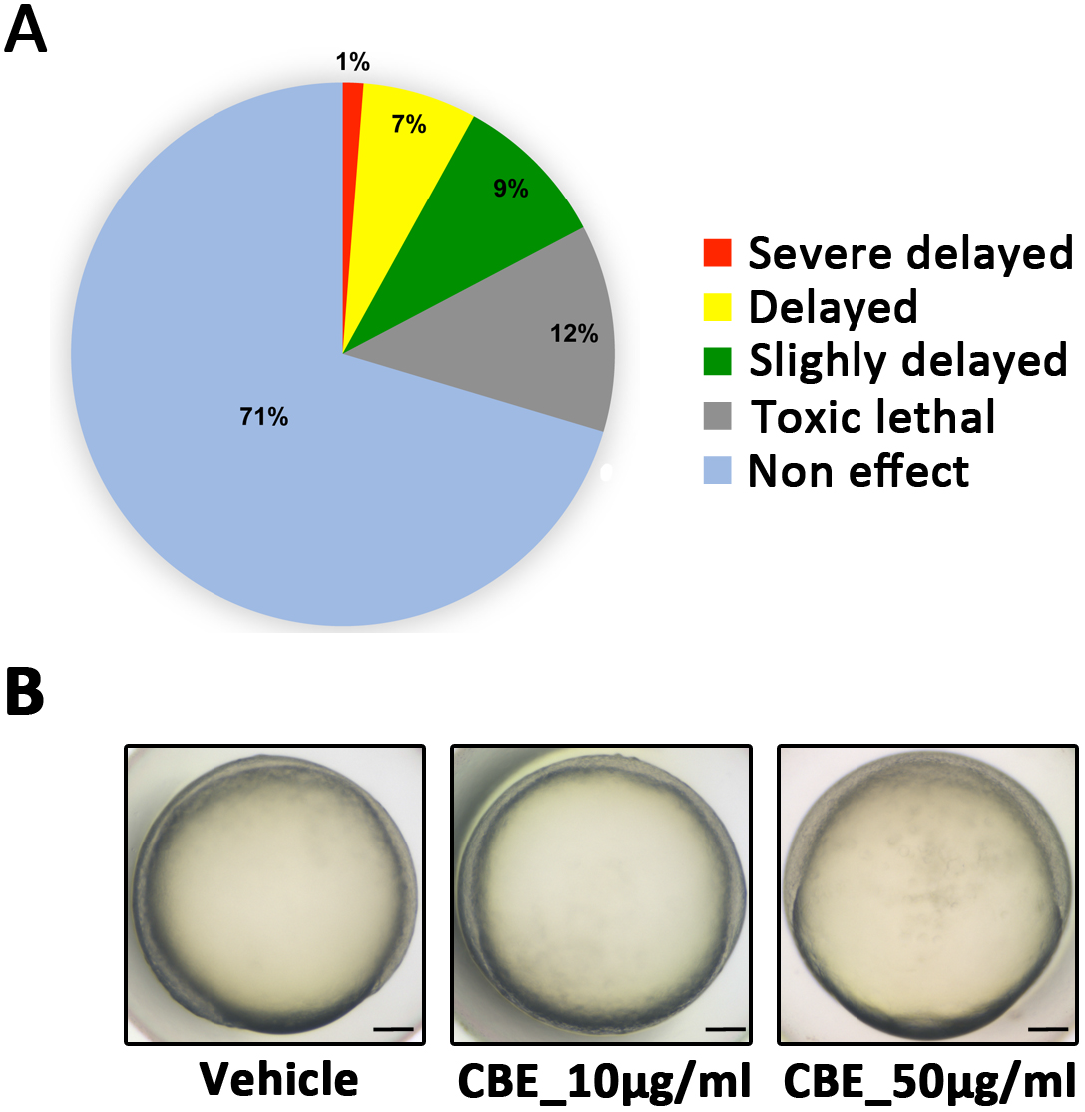
Zebrafish embryo screen identified CBE as an epiboly-interrupting herbal medicine. (A) Cumulative results of the phenotypic screen in which each herbal medicine was tested at 10 μg/ml. A total of 162 herbal medicines were tested in the screening assay. A positive “hit” was defined as an herbal medicine that interfered with epiboly progression. (B) Representative images of embryos treated with vehicle (left), 10 μg/ml CBE (middle), or 50 μg/ml CBE (right). Scale bar, 100 μm.

### CBE suppressed metastatic dissemination of human cancer cells in a zebrafish xenotransplantation model

We examined whether CBE could suppress cell motility and invasion of human cancer cells through a Boyden chamber assay. Before conducting the experiment, we investigated whether the extract might affect viabilities of highly metastatic human cancer cell line, MDA-MB-231 cells. The assay showed that CBE did not affect the proliferation of the cells (Figure 2A). A Boyden chamber assay revealed that CBE inhibited cell motility and invasion of MDA-MB-231 cells in a dose-dependent manner (Figure 2B). Furthermore, we examined whether CBE could suppress metastatic dissemination of the cells in a zebrafish xenotransplantation model. RFP-labeled MDA-MB-231 (231R) cells were injected into the duct of Cuvier of *Tg (kdrl: eGFP)* zebrafish at 2 dpf and then maintained in the presence of either vehicle or CBE. Twenty-four hours post-injection, the numbers of fish showing metastatic dissemination of 231R cells were measured via fluorescence microscopy. Three independent experiments revealed that the frequencies of fish in CBE-treated group showing head, trunk, or end-tail dissemination, significantly decreased to 30.7±0.3%, 13.5±4.4% or 49.1±1.2% when compared with those in the vehicle-treated group; 62.7±5.4%, 36.4±23.2% or 93.3±9.4%. Conversely, the frequency of the fish in CB-treated group not showing any dissemination, significantly increased to 49.4±3.1% when compared with those in the vehicle-treated group; 3.3±4.7% (Figure 2C and D). These results indicated that CBE suppressed metastatic dissemination of human cancer cells *in vivo*.

**Figure 2.**
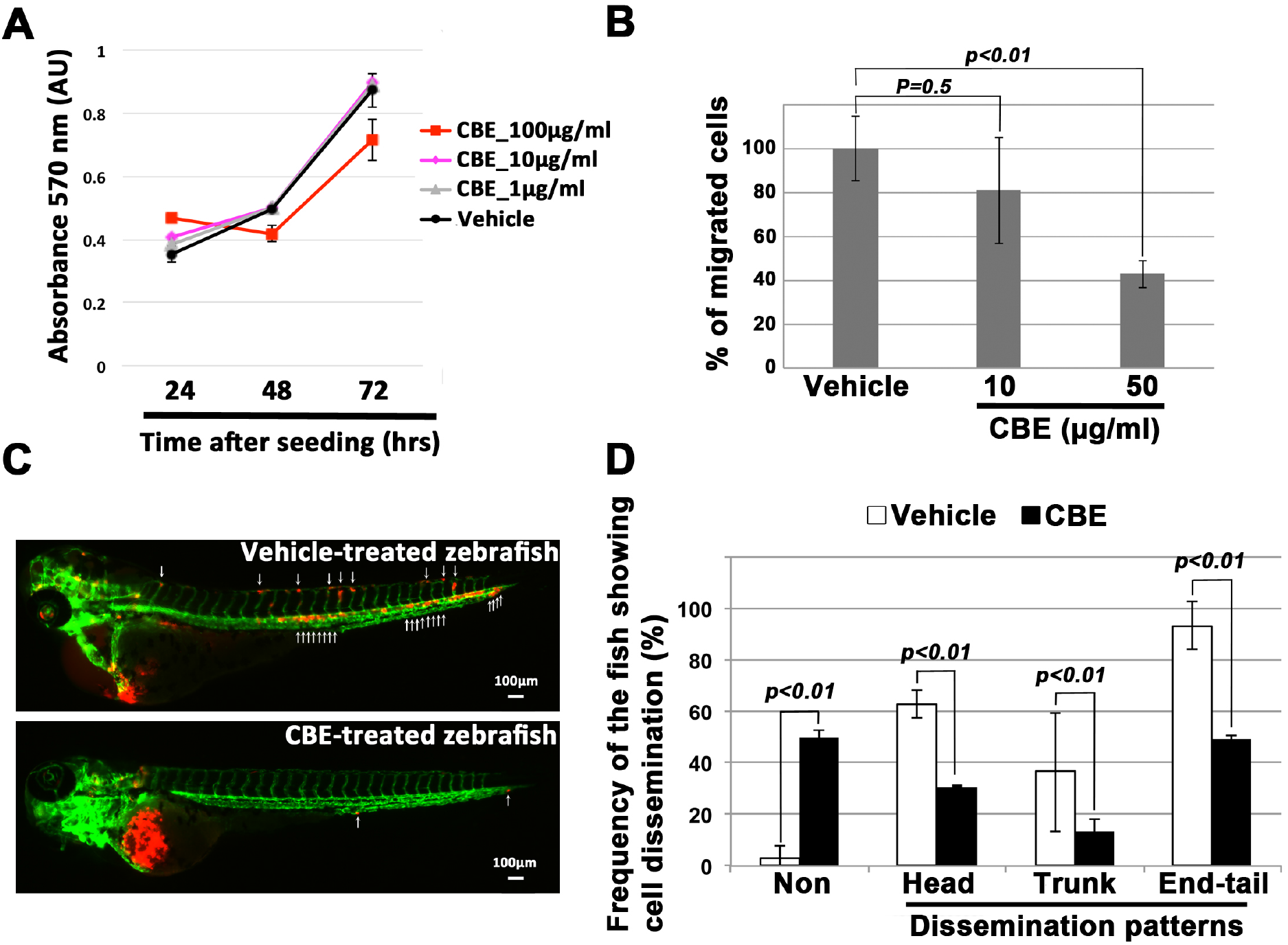
CBE suppressed metastatic dissemination of human cancer cells *in vitro* and *in vivo*. (A) Effect of CBE on proliferation of MBA-MB-231 cells using the MTT assay. (B) Effect of CBE on cell motility and invasion of MBA-MB-231 cells. The cells were treated with vehicle or CBE and then subjected to the Boyden chamber assay. 1% fetal bovine serum (FBS) (v/v) was used as the chemoattractant in both assays. Each experiment was performed at least twice. (C) Representative images of dissemination of RFP-labeled MDA-MB-231 (231R) cells in zebrafish. The zebrafish inoculated with 231R cells were treated with either vehicle (top panel) or CBE (bottom panel). White arrowheads indicate disseminated 231R cells. The images are shown at 4× magnification. Scale bar, 100 μm. (D) The mean proportions of zebrafish with head, trunk, or end-tail dissemination are presented in a graph. Each value is provided as the mean ± SEM of two independent experiments. Statistical analysis, Student’s t test.

### CBE inhibited glycolytic metabolism

Finally, we elucidated the mechanism of action of how CB-extract suppressed metastatic dissemination of cancer cells. Recent study revealed that genetic inhibition of MondoA protein which shows transcriptional activities in presence of glucose likely glucose-6-phosphate (G6P), interrupts epiboly progression in zebrafish embryos [11]. Moreover, other studies demonstrate that cinnamon reduces blood glucose levels in patients with type 2 diabetes [12]. Thus, we hypothesized that CBE might impair glycolytic metabolism in cancer cells. Extracellular acidification rate (ECAR) analysis showed that the maximum cellular glycolytic capacity was lower in CBE-treated MDA-MB-231 cells than in vehicle-treated the cells (Figure 3A). Furthermore, quantitative metabolome analyses using a gas chromatography mass spectrometry (GC-MS) revealed the glucose level was six times higher in CBE-treated MDA-MB-231 cells than in vehicle-treated the cells. Conversely, the level of G6P, which is the phosphorylation product of glucose, was 50% lower in CBE-treated MDA-MB-231 cells than in vehicle-treated cells. No fructose 6-phosphate (F6P), which is produced by isomerization of G6P, was detected in CBE-treated MDA-MB-231 cells (Figure 3B). Furthermore, qPCR and western blot analyses revealed that hexokinase 2 (HK2) expression was lower in CBE-treated cells than in vehicle-treated the cells (Figure 3C). These results indicated CBE inhibited glycolytic metabolism in MDA-MB-231 cells.

**Figure 3.**
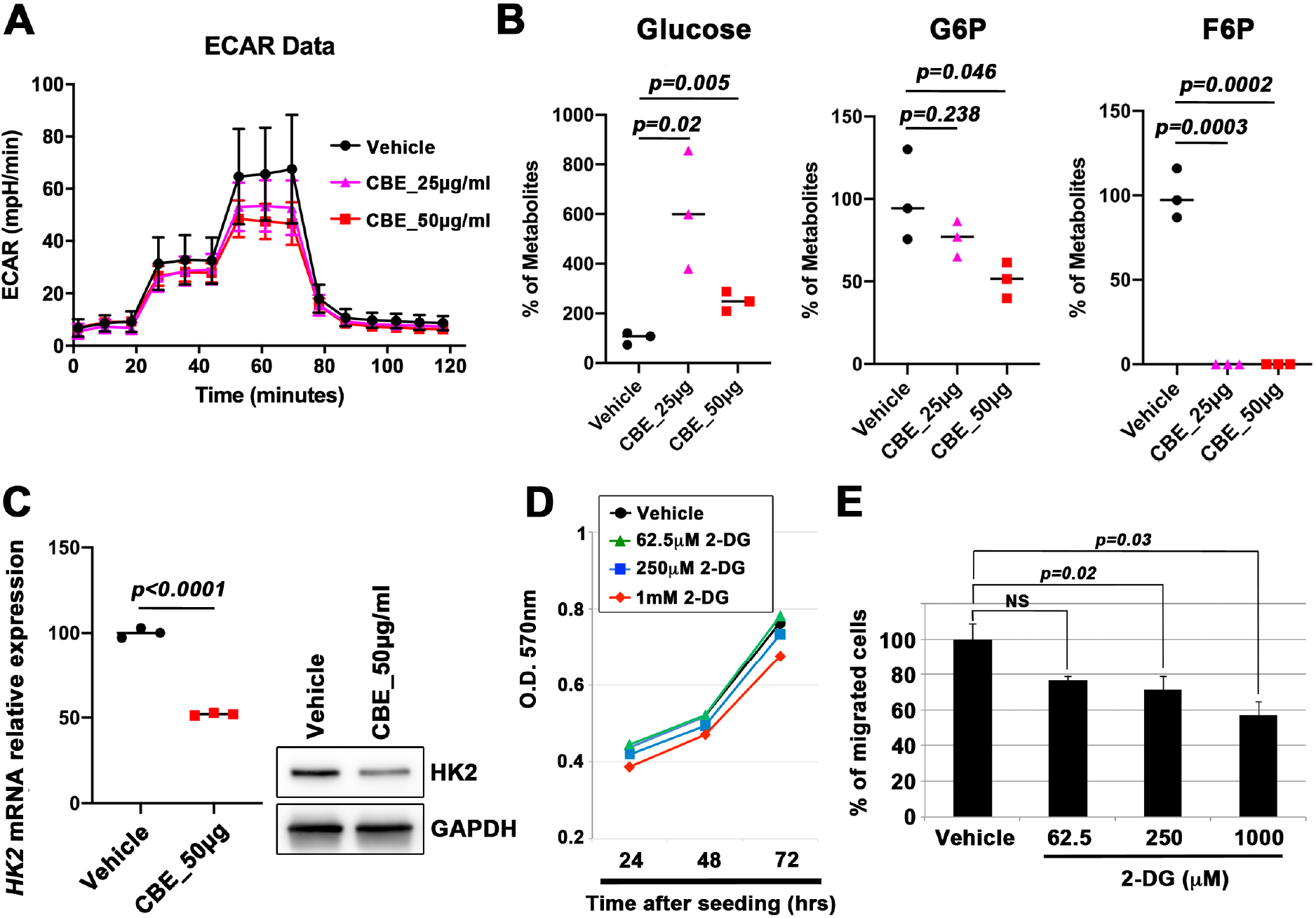
CBE inhibited glycolytic metabolism. (A)Representative glycolytic rate profile of either vehicle- or CBE-treated MDA-MB-231 cells. MDA-MB-231 cells were cultured in the presence of vehicle, 25 μg CBE, or 50 μg CBE for 6 h before measuring the extracellular acidification rate (ECAR) using Seahorse XF. Cellular levels of glucose, glucose 6-phosphate, and fructose 6-phosphate were measured by GC-MS. (C) HK2 expression levels in vehicle- or CBE-treated MDA-MB-231 cells were assessed by qPCR (left) and western blot (right) analyses. (D) Effect of 2-deoxy-D-glucose (2-DG) on proliferation of MDA-MB-231 cells was measured by MTT assay. (E) Effect of 2-deoxy-D-glucose (2-DG) on cell invasion and migration of MDA-MB-231 cells was measured by the Boyden chamber assay.

We confirmed whether pharmacological inhibition of HK2 with 2-deoxy-D-glucose (2-DG), which is a glucose analog that acts as a competitive inhibitor of glucose metabolism, might show same effect. Treatment with 2-DG suppressed cell motility and invasion of MDA-MB-231 cells without affecting cell viability (Figure 3D and E). These results showed that blocking the HK2 activity was sufficient to suppress cell invasion and migration of the cells. We concluded that CBE suppressed metastatic dissemination of cancer cells via inhibition of glycolytic metabolism.

## Discussion

CB is one of traditional herbal medicines and includes several bio-active compounds: cinnamic aldehyde, cinnamyl aldehyde, carbohydrate, tannin and mucus. Our study demonstrated that CBE suppressed metastatic dissemination of human cancer cells through downregulating glycolytic metabolism. Although we have not addressed which CBE compound has this suppressor effect, we propose that daily CB consumption may have potential to suppress cancer progression.

We demonstrated that CBE decreased HK2 expression and that disrupted the production of G6P and F6P, which are intermediate metabolites of glycolytic metabolism. Recent studies report that HK2 promotes metastasis [13, 14]. Therefore, HK2 represents a potential target for suppressing metastasis progression. Hence, we predict that the identification of the CBE compound causing the downregulation of the HK2 expression may lead to the discovery of an anti-metastatic drug.

## Acknowledgments

We sincerely appreciate Prof. Hayakawa (Institute of natural medicine, University of Toyama) for providing the herbal medicine library to us. We thank Dr. Shimada (Mie University) for teaching methods of zebrafish xenotransplantation to us. This research was performed as a part of the Cooperative Research Project with Institute of Natural Medicine, University of Toyama in 2021. This work was supported in part by research funds from the Yamagata prefectural government and the City of Tsuruoka.

